# Unbiased transcriptomic analysis of chondrocyte differentiation in a high-density cell culture model

**DOI:** 10.1101/2021.03.20.436275

**Authors:** Claudia Kruger, Aimee Limpach, Claudia Kappen

## Abstract

In the developing vertebrate skeleton, cartilage is an important precursor to the formation of bones. Cartilage is produced by chondrocytes, which derive from embryonic mesoderm and undergo a stereotypical program of differentiation and maturation. Here we modeled this process *in vitro*, using primary fetal mouse rib chondrocytes in a high-density cell culture model of cartilage differentiation, and performed genome-wide gene expression profiling over the course of culture. The overarching goal of this study was to characterize the molecular pathways involved in cartilage differentiation and maturation. Our results also enable a comprehensive appraisal of distinctions between common *in vitro* models for cartilage differentiation, and of differences in their molecular resemblance to cartilage formation *in vivo*.

## INTRODUCTION

Most of the bones in the vertebrate body are formed through the process known as endochondral ossification. Progenitor cells for this process derive from embryonic mesoderm, specifically from paraxial and lateral mesoderm. These precursors assemble into so-called condensations, with greater cell density than the surrounding mesenchyme^1^. Within these condensations, cells commit to chondrocyte lineage differentiation, largely under control of the transcription factor Sox9^2, 3^, while continuing to proliferate as the structures that will make up the future skeleton grow with increasing age of the embryo. Eventually, cells exit the cell cycle and enlarge their cytoplasms, and through action of the transcription factor Runx2^4–6^ mature into hypertrophic chondrocytes. The components and composition of extracellular matrix secreted by the cells change with time, and eventually become mineralized and ossified^7^.

*In vitro* modeling of this process requires special culture conditions, because cell-matrix interactions as well as cell-cell contacts are needed for chondrocytes to differentiate, mature, and maintain the chondrocyte phenotype. Micromass cultures of embryonic limb mesenchyme^8, 9^ as well as high-density cultures of rib chondrocytes have been used by us^10, 11^ and others^12–16^ to investigate molecular mechanisms in cartilage differentiation. Many of these studies focused on the roles of specific individual regulators or signaling pathways, as reviewed by Liu et al., 2017^17^. In order to provide a systematic survey of molecular pathways critical to chondrocyte differentiation, maturation and function, we conducted unbiased genome-wide gene expression profiling on primary chondrocytes isolated from fetal mice and differentiated in culture. Our results reveal changing expression repertoires over time in culture, with overall reduction in the number of gene expressed, and progressive activation of gene subsets that define the molecular phenotype of mature chondrocytes.

## METHODS

### Preparation of mouse rib chondrocytes

All work with animals was approved by the Institutional Animal Care and Use Committee in accordance with the GUIDE FOR THE CARE AND USE OF LABORATORY ANIMALS (https://grants.nih.gov/grants/olaw/guide-for-the-care-and-use-of-laboratory-animals.pdf). Pregnant FVB mice were sacrificed at day E18.5 of gestation and fetal rib cages were prepared as described by Cormier et al.^10^. Enzymatic digestion with Collagenase (3mg/ml, Sigma) and 0.25% Trypsin/EDTA (Invitrogen/Gibco BRL) in PBS was performed for 45 minutes at 37°C. Further digestion and preparation of cells followed our previously published method^10^.

### Primary chondrocyte high density cultures

Cells were plated on 35mm dishes coated with 0.1% gelatin at a final concentration of 1×10^6^cells/well in DMEM/F12 (Gibco BRL), high glucose, 10% fetal bovine serum and 1% Penicillin/Streptomycin antibiotics. The medium was supplemented with 25mg/ml Ascorbic Acid (Sigma) and 10mM *β*-glycerolphosphate to differentiate chondrocytes^10^. Cultures were incubated in a 37°C, 5% CO_2_ incubator for up to 21 days, with changes of culture medium every other day.

### Cell staining

To assess cartilage formation and chondrocyte cellular phenotype, cells were fixed in 70% ethanol for 30 minutes and then stained with either Alcian Blue (15mg Alcian Blue Stain; 30ml 95% Ethanol and 20ml Glacial acetic acid) or Alizarin Red (1g Alizarin Red; 50ml distilled water; 1% KOH) for 1 hour at room temperature. After rinsing with 70% ethanol, cells were dehydrated through an ethanol series before cover slipping on the dish.

### Immunohistochemistry

The production of Collagen II was assessed by immunocytochemistry using a mouse monoclonal antibody (Chemicon), appropriate HRP-conjugated secondary antibody (Invitrogen) and DAB colorimetric reaction (Sigma). Type X collagen deposition in the extracellular matrix was analyzed by immunocytochemistry using a mouse monoclonal antibody (RDI) and the same secondary reagents as described above.

### RNA isolation and Microarrays

For the microarray studies, RNA was isolated from cultured chondrocytes at culture days 2, 6, 9, 12, and 15 as follows: cells were scraped off 35mm dishes in 0.5ml Trizol reagent (Invitrogen). Chloroform was added (20% of volume), followed by vortexing and incubation at room temperature for 5 minutes. After 10 minutes of 10,000rpm centrifugation at 4°C, the aquaeous layer was precipitated with 250ml of isopropanol, vortexed and incubated at room temperature for 10 minutes, followed by 10 minutes of 13,000rpm centrifugation at 4°C. The pellet was washed twice with 75% ethanol, then briefly air dried. RNA was solved in nanopure DEPC-treated water, concentration was measured spectrophotometrically, and quality was confirmed with an Agilent 2100 Bioanalyzer. Preparation and hybridization of microarrays was conducted according to the Affymetrix Expression Analysis Technical Manual using Mouse Genome 430 2.0 Microarray Chips. Arrays were scanned using a Gene-Chip 3000 scanner, with averaging of signals from two sequential scans. The data are accessible in the Gene Expression Omnibus (GSE102292).

### Microarray data analysis

Chip data were analyzed using the Microarray Suite (MAS) 5.0 (Affymetrix) which assigned an expression call (P=present, M=marginal, A=absent), based on comparison of the matched and mismatched probes for each respective gene. Results were normalized for each chip using the Global Scaling Method (Affymetrix). CHP files were imported into GeneSpring GX10 software (Silicon Genetics), with default transformation.

K-Means clustering was performed using JMP Genomics software 5.1 built on JMP9 and SAS9.3 (SAS Institute Inc.) using normalized values from the CHP file with log2 transformation. Probe IDs with intensity signals below 500 at all time points were eliminated. 12078 probe sets with values above 500 in at least one out of five time points were sorted ascending based on culture day 2, split into quartiles and median normalized. Three clusters per quartile were obtained to present increasing, decreasing and unchanged intensity values over time. A total of 12 clusters were subjected to the functional annotation tool in DAVID (http://david.abcc.ncifcrf.gov) with default of medium stringency, followed by confirmation of major categories with increasing stringency level.

Expression profiles of cultured cells were compared to freshly isolated cells using the microarray data for primary chondrocytes from 8 individual control mice published in our earlier paper (Kruger and Kappen, 2010^18^; GEO accession number GSE19002) Functional annotation was performed in DAVID as described above, supplemented with information from Mouse Genome Informatics (http://www.informatics.jax.org/). The Riken Transcription Factor Database (http://genome.gsc.riken.jp/TFdb) was used to obtain 1675 mouse transcription factor genes.

Comparison of our datasets to the microarray data published for micromass cultures of embryonic limb mesenchymal cells in cartilage differentiation conditions (James et al., 2005^19^; GEO accession number GSE2154) and to gene expression profiling in dissected zones from embryonic tibiae (James et al., 2010^20^; GEO accession number GSE7685) was possible because those surveys also used Affymetrix platforms. Based on probe identifiers represented on the oldest arrays (James et al., 2005^19^), we constructed a dataset for all probes that displayed expression, yielding 11529 informative probes. All data were in log-transformed format, and analyzed in CyberT and JMP Genomics; Principal Component Analysis used using JMP default settings.

### Primers

Primers for amplification were designed using Primer Express 3 software (Applied Biosystems) described in Kruger et al. 2006^21^. In brief: TM requirements: min. TM 58°C, max. TM 60°C, optimal TM 59°C; GC content requirements: min. % GC 20, max. % GC 80; length requirements: min. length 9, max. length 40; optimal length 20; amplicon requirements: min. TM 0°C, max. TM 85°C, min. length 50, max. length 150. Primers were designed to span an exon–exon junction, where possible; primer sequences are listed in Supplemental Table 1.

### Quantitative real-time PCR

Reverse transcription of RNA was performed using SuperScript III First-Strand Synthesis System (Invitrogen), as described in Kruger and Kappen, 2010^22^. Purification of cDNA used the Qiagen PCR purification kit, and cDNA concentration was measured on the NanoDrop ND-1000 Spectrophotometer. Each PCR reaction (25µl) was performed on the cDNA equivalent of 4ng of RNA, in triplicates, with iTaq SYBR Green Supermix with ROX (Bio-Rad Laboratories) using the ABI PRISM 7900HT Sequence Detection System (Applied Biosystems)^22^.

Data analysis used the SDS software version 2.2.2 (Applied Biosystems) with default baseline of 3 to 15 cycles and a threshold value of 0.04. Triplicate measurements for a given sample were averaged and normalized to measurements for Gapdh cDNA in the same sample. Gapdh was chosen as the reference to provide comparability with our prior cartilage gene expression analyses^21, 22^. Relative expression compared to day 2 was calculated using the Comparative C_T_ method: 2^-^*^ΔΔ^*^C^_T_, where *ΔΔ*C_T_ = *Δ*Ct_day6,9,12,15_ – *Δ*Ct_day2_. Average amplification efficiencies (Supplemental Table 1) were determined for each gene-specific reaction over the first three cycles above threshold: *Δ*Rn_cycle (n) / *Δ*Rn_cycle (n-1) and replaced the value of 2 in 2^-^*^ΔΔ^*^C^_T_ in order to calculate the actual fold-change of expression.

## RESULTS

Over the course of culture *in vitro* under conditions of high cell density, primary fetal rib chondrocytes differentiate from their initial small size into hypertrophic cells (Figure 1). Alcian blue (Panel A) staining identifies cartilage-producing cells by 6-9 days in culture, and by day 15, almost all cells are producing polysaccharides that react with the dye. In late-stage cultures, the extracellular matrix becomes mineralized, as indicated by Alizarin Red staining (Panel B). Collagen II positivity is found in most cells in early cultures, and is retained by fewer cells at later stages (Panel C). Collagen X staining identifies hypertrophic chondrocytes, particular in later stage cultures (Panel D). Thus, primary neonatal mouse rib chondrocytes in high-density culture recapitulate the known differentiation program of endochondral cartilage.

**Legend to Figure 1.**
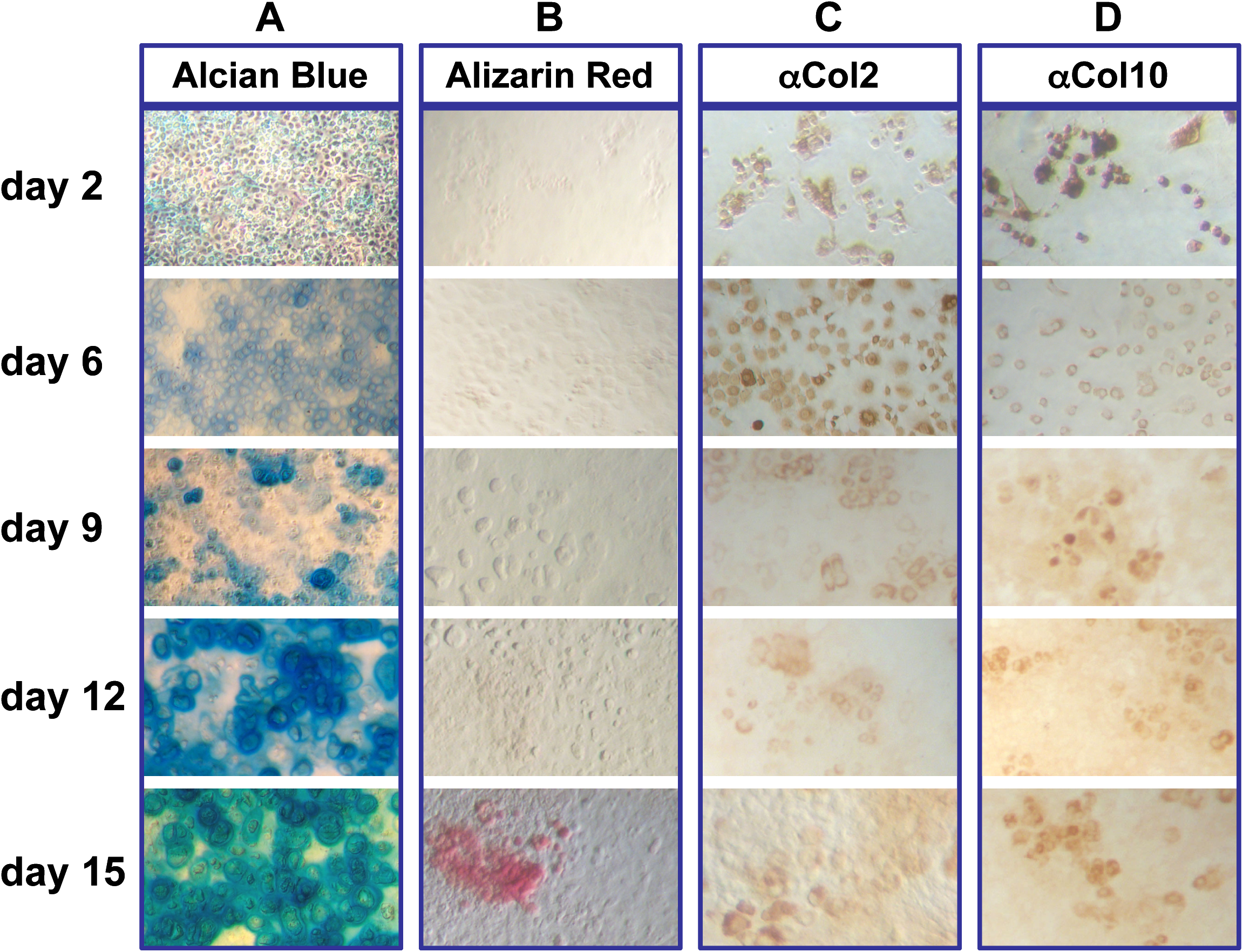
High density chondrocyte culture system. Representative primary chondrocyte cultures plated at 1 x 10^6^ cells/well in 6-well plates and cultured for the number of days indicated. Morphology of chondrocyte cultures was examined using light microscopy (20X) of cells stained with **A**) Alcian blue, **B**) Alizarin red, **C**) Collagen 2 and **D**) Collagen 10.

In order to obtain an unbiased survey of the molecular mechanisms involved in this stereotypic differentiation program, we conducted microarray analyses at various time points of culture, starting from day 2. This time point was chosen to avoid contamination from dying cells that did not accommodate to the culture conditions. To account for temporal dependency, we analyzed the prior relative to the following samples at each time point, so that the day 2 profile was compared to all later time points combined (days 6, 9, 12, 15), day 2 and 6 profiles together were compared to the remaining later stages (day 9, 12, 15), and so on, as outlined in Figure 2.

**Legend to Figure 2.**
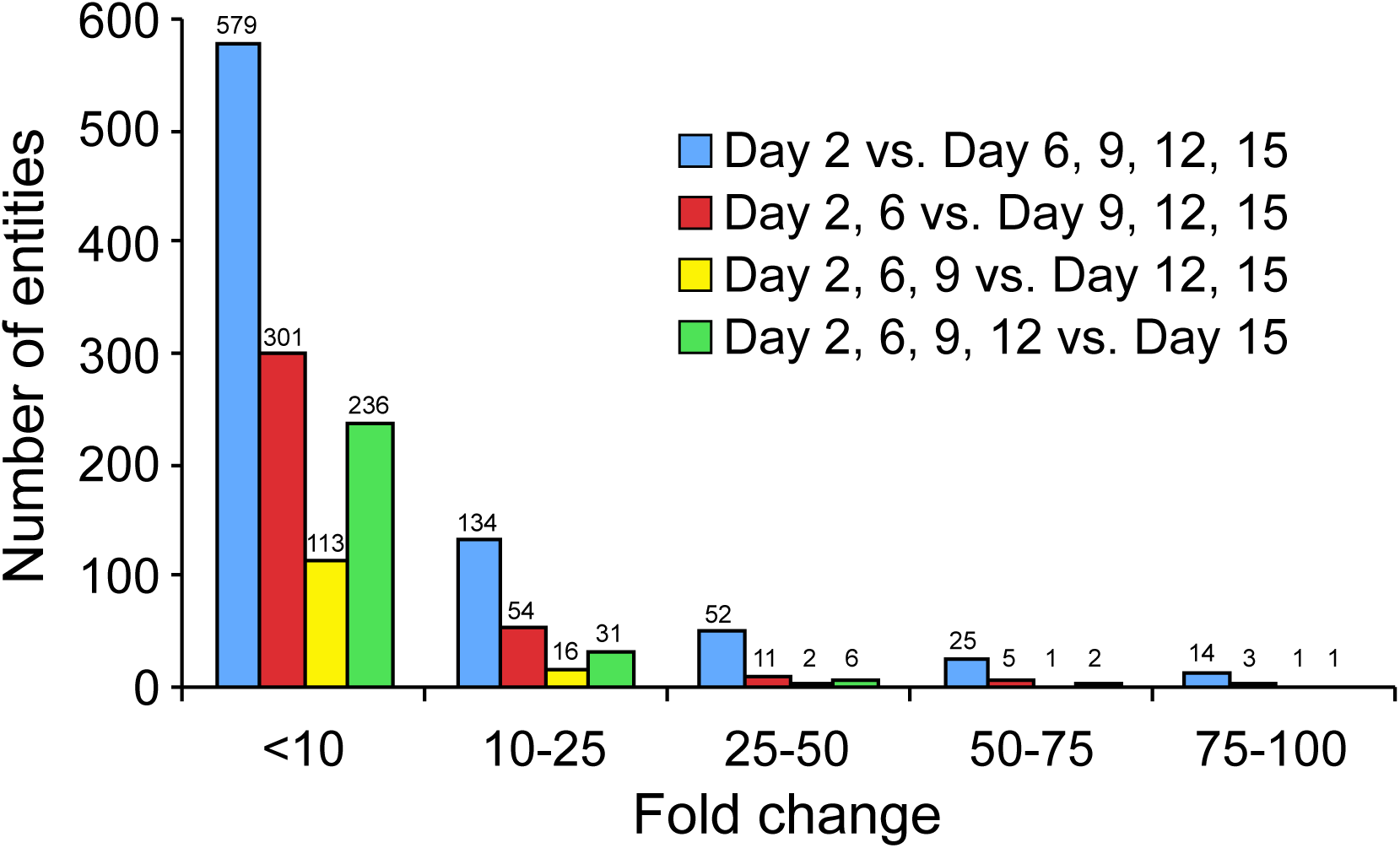
Distribution of differential expressed genes in high density chondrocyte culture systems over time. Affymetrix Mouse Genome 430 2.0 Microarray Chips were probed with RNA obtained from chondrocytes after 2, 6, 9, 12 and 15 days in culture. Microarray data were grouped like indicated and analyzed in GeneSpring. Graphed are the number of entities with a “present flag” in at least 1 out of 5 samples and a fold-change greater than 10.

The greatest difference in any of these analyses was found between early and later time points, with over 800 entities being expressed at different levels on day 2 compared to days 6, 9, 12 and 15 combined. Levels of gene expression differed from below 10-fold (72% of the entities), up to 100-fold and over (1.7% of the entities). The comparisons at intermediate time points of culture identified fewer differentially expressed entities, and very few genes with large fold-changes. However, the latest stage of culture profiled, day 15, again exhibited noticeable differential gene expression, with 276 entities passing the significance criteria. These results indicate that chondrocytes undergo greater molecular changes early in the culture, followed by maturation and relatively moderate changes at later stages.

This interpretation is supported by cluster analyses of those entities that change over the entire course in culture (Figure 3). The top 6 (by gene number) clusters are depicted in the heat maps, with curves for individual entities in each cluster given below. This analysis reveals the existence of multiple distinct temporal patterns, with expression levels predominantly increasing from early stages of culture: compared to 479 entities in clusters 1, 2, 4-6 (82.7%), only 100 entities exhibited declining levels in cluster 3 (17.3%). Greater fractions of entities with declining expression are seen after 6 days in culture, and by days 9 and 12 (23%, 46% and 83.5%, respectively), accompanied by progressively smaller fractions of entities with increasing expression (76.9%, 54%, and 16.5%, respectively). Furthermore, the inflection points of curves indicate that time points of greatest change differ for various clusters, revealing distinct temporal trajectories for particular groups of genes (top pathways in each cluster are given in Supplemental Table 2, with transcription factors listed in Supplemental Table 3).

**Legend to Figure 3.**
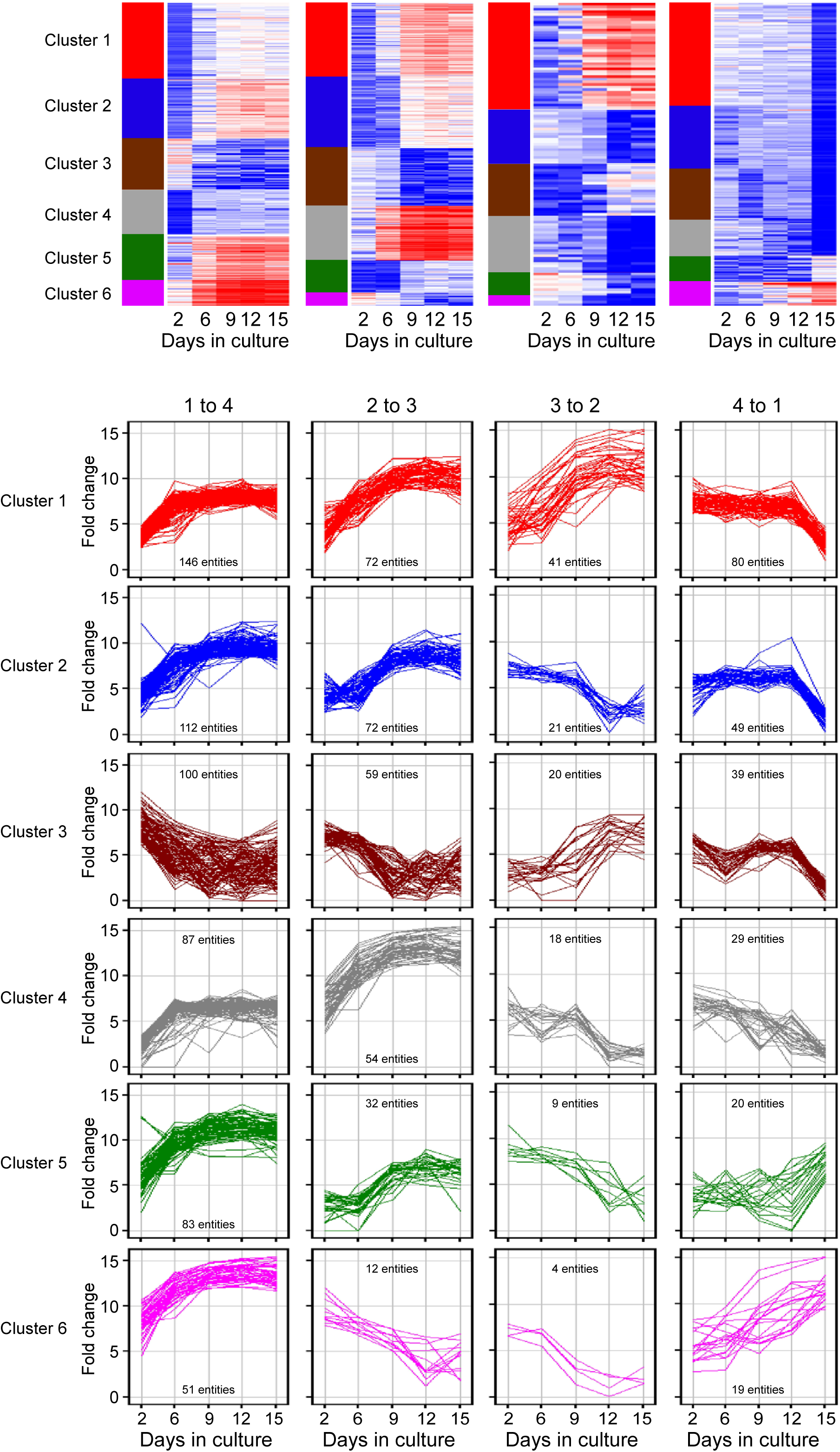
Time series microarrays of chondrocyte differentiation in culture. Primary chondrocytes were seeded into high density cultures. RNA was isolated at days 2, 6, 9, 12 and 15 and processed for hybridization to Affymetrix oligonucleotide arrays 430 2.0. MAS 5.0 software was used for normalization after which the data were transferred to GeneSpring for further analysis. Samples were grouped like indicated. Entities which passed the criteria “present flag in at least 1 out of 5 samples and an expression level more than 10-fold” underwent K-means clustering with 50 iterations. For each comparison identified genes were grouped in 6 clusters.

In order to validate the results with an independent method, we performed quantitative real-time PCR (qRT-PCR) assays. For each cluster in Figure 3, we designed primers for genes known to play a role in cartilage and bone development, such as for example, Collagens and Bmp4. Figure 4 shows that the expression levels measured by qRT-PCR are consistent with the temporal pattern of the respective cluster in the microarray analyses. Notably, vast differences were found in the magnitudes of change, with transcript levels of Col4a6 changing moderately by 5-fold, whereas mRNA levels of the Otor gene that encodes a protein closely related to cartilage-derived-retinoic-acid-sensitive-protein (CD-RAP/MIA) change up to 1000-fold. The large increase in Otor gene transcription is likely functionally involved in cartilage maturation, as it has previously been reported that Otor antisense treatment of mesenchymal cells decreases chondrogenesis in micromass cultures (Cohen-Salmon, 2000)^23^. Similarly, hedgehog-interacting-protein (Hhip) transcript levels increase with time, indicating that hedgehog signaling and the Hhip-mediated feedback control is active in our cultures^24^. Consistent with progressive cartilage maturation, expression levels of collagen genes and metalloproteinases increase over time, while Sox11 and Dlx1 levels decrease as cells cease to proliferate. Thus, the temporal profiles of numerous genes during culture provide strong evidence for the fidelity of high-density cultures as models of endochondral cartilage formation and maturation.

**Legend to Figure 4.**
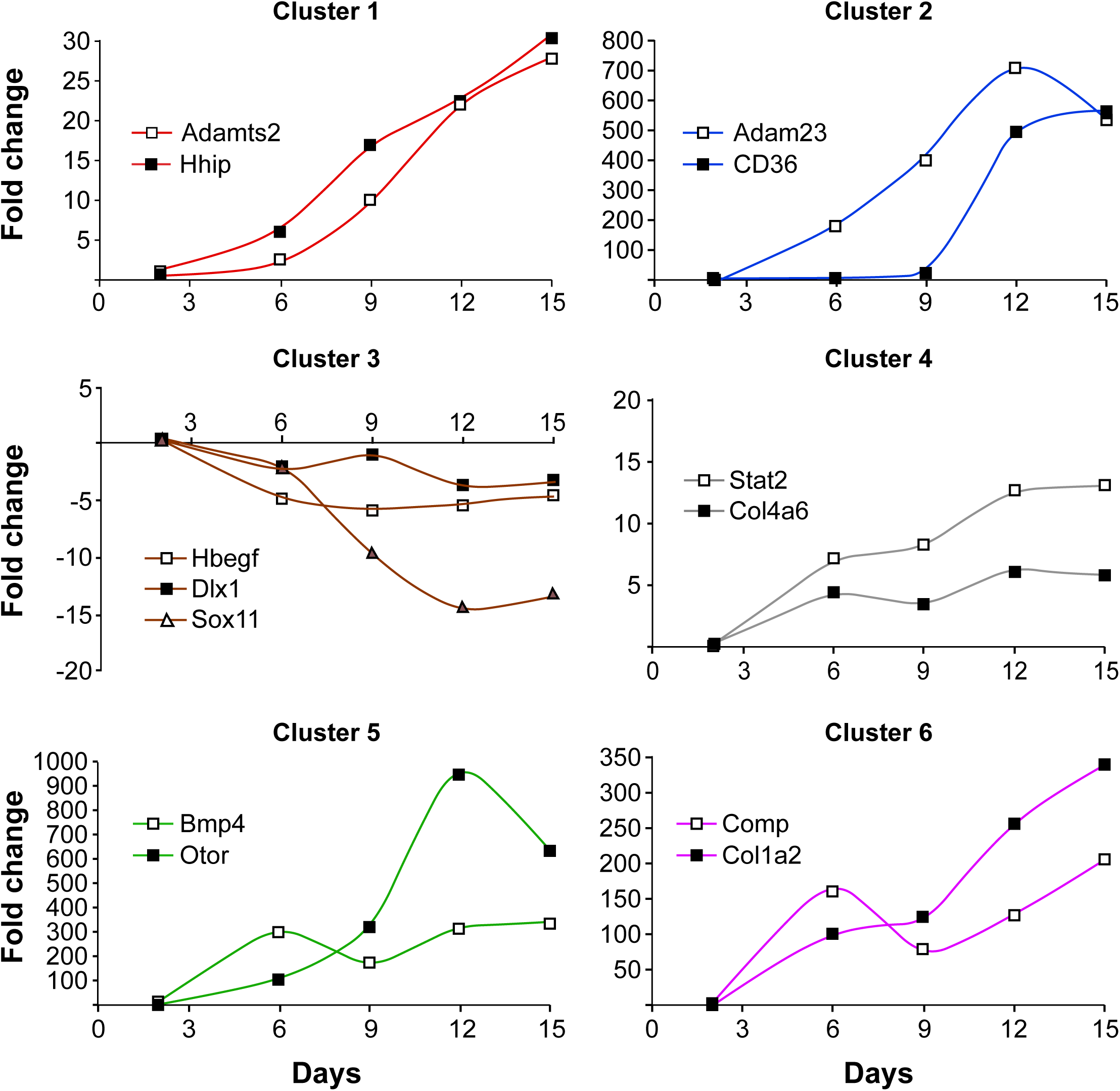
Validation of microarray analyses by quantitative RT-PCR. Candidate genes were selected from the microarray comparison day 2 versus all other time points. Three genes were selected from cluster 3 for the validation by qRT-PCR and two genes from the respective remaining clusters. Individual samples were run in triplicates and normalized to expression levels of Gapdh. The fold-change was calculated using the formula 2^-^*^ΔΔ^*^C^_T_, with the amplification efficiency for each gene accounted for. All time points were compared to day 2.

We then wanted to assess, from a genome-wide perspective, the extent to which cultured cells resemble primary chondrocytes at the time of isolation. To this end, we compared the gene expression profile of day 2 cultures to profiles of freshly isolated chondrocytes from neonatal mouse ribs^18^. Intriguingly, substantial differences were found (Figure 5): 4544 entities displayed expression differences of over 2-fold, 879 entities differed more than 5-fold, and 78 entities by over 25-fold. Among the transcripts with stronger representation in primary rib cartilage samples were those encoding Hemoglobin components, suggesting that these fresh cell preparations might have contained some erythrocytes. Expression of Erythroid-associated-factor (Eraf, also known as *α*-hemoglobin-stabilizing-protein Ahsp) and Erythroid-aminolevulinic-acid-synthase-2 (Aras2) in fresh cell mRNA is consistent with this interpretation. Also over-represented in the primary cell preparation were transcripts encoding Myosin-light-chain, Myogenic-factor-5, Myogenin, and other muscle markers, indicating that some cells from intercostal muscles were present. Thus, expression profiles of freshly isolated primary rib cartilage reflect the *in vivo* source of the chondrocytes, with some admixture by several other identifiable cell types.

**Legend to Figure 5.**
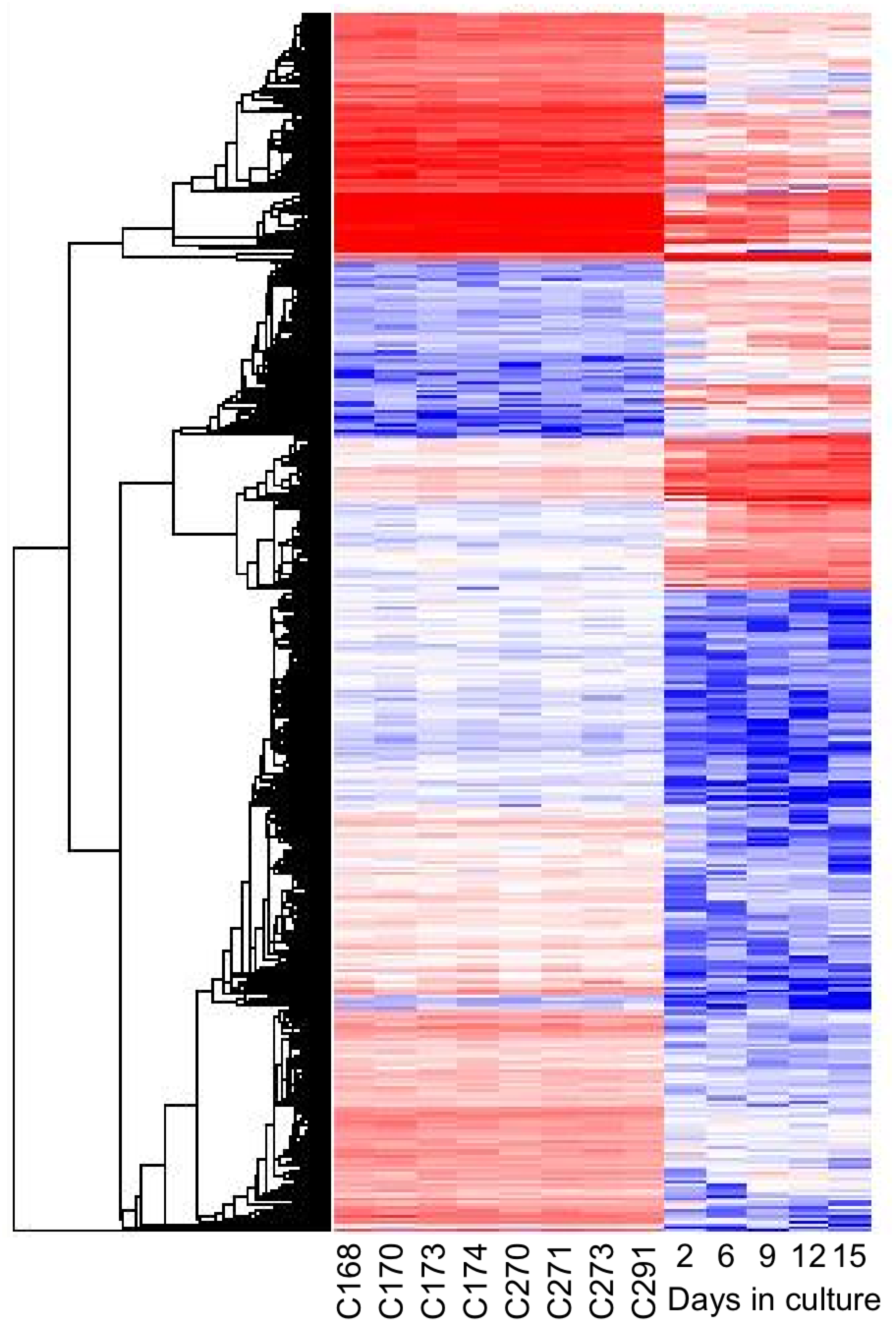
Gene expression profiles of normal and cultured chondrocytes. Graphic representation of relative expression levels after hierarchical cluster analysis, each line represents one gene. Cultured chondrocytes were harvested at day 2, 6, 9, 12, 15 and gene expression was compared to primary chondrocytes obtained from four Hoxc8 and four Hoxd4 control animals at gestational day 18.5. Data mining was performed with GeneSpring (baseline transformation), samples were averaged over replications in each condition. In an initial step 23474 entities passed the criterion “present flag” in at least 5 out of 13 samples. Statistical analysis was performed using the unpaired t-Test with unequal variance and no further correction. A total of 879 entities were identified with expression levels that were changed in cultured chondrocytes by more than 5-fold (P<0.05) compared to primary chondrocytes.

Generally, day 2 cultured chondrocytes exhibited lower expression for over 70% of the differentially represented transcripts and gained higher expression for only a minor fraction of genes, among them transcripts encoding proteins involved in oxidation-reduction processes, lipid metabolism and mitochondrial function. Examples are NAD-kinase (Nadk), Glutaredoxin (Glrx), Glutathione-peroxidase (Gpx4), Carnitine-palmitoyltransferases (Cpt1a, Cpt2), Lysosomal-acid-lipase (Lipa), Fatty-acyl-Coenzyme-A-reductase (Fra), and Acyl-Coenzyme-A-acyltransferases (Acaa1b, Acaa2). However, the vast majority of genes passing the statistical significance test in the fresh-vs.-cultured comparison exhibited modest differences of expression levels, in the range of 1.3-1.4 fold (Figure 6). Given that the dataset for fresh chondrocytes was generated in a separate experiment, with RNA isolations then^18^ and microarray assays here performed by different experimenters, the biological significance of moderate changes in expression level revealed by these analyses remains to be investigated.

**Legend to Figure 6.**
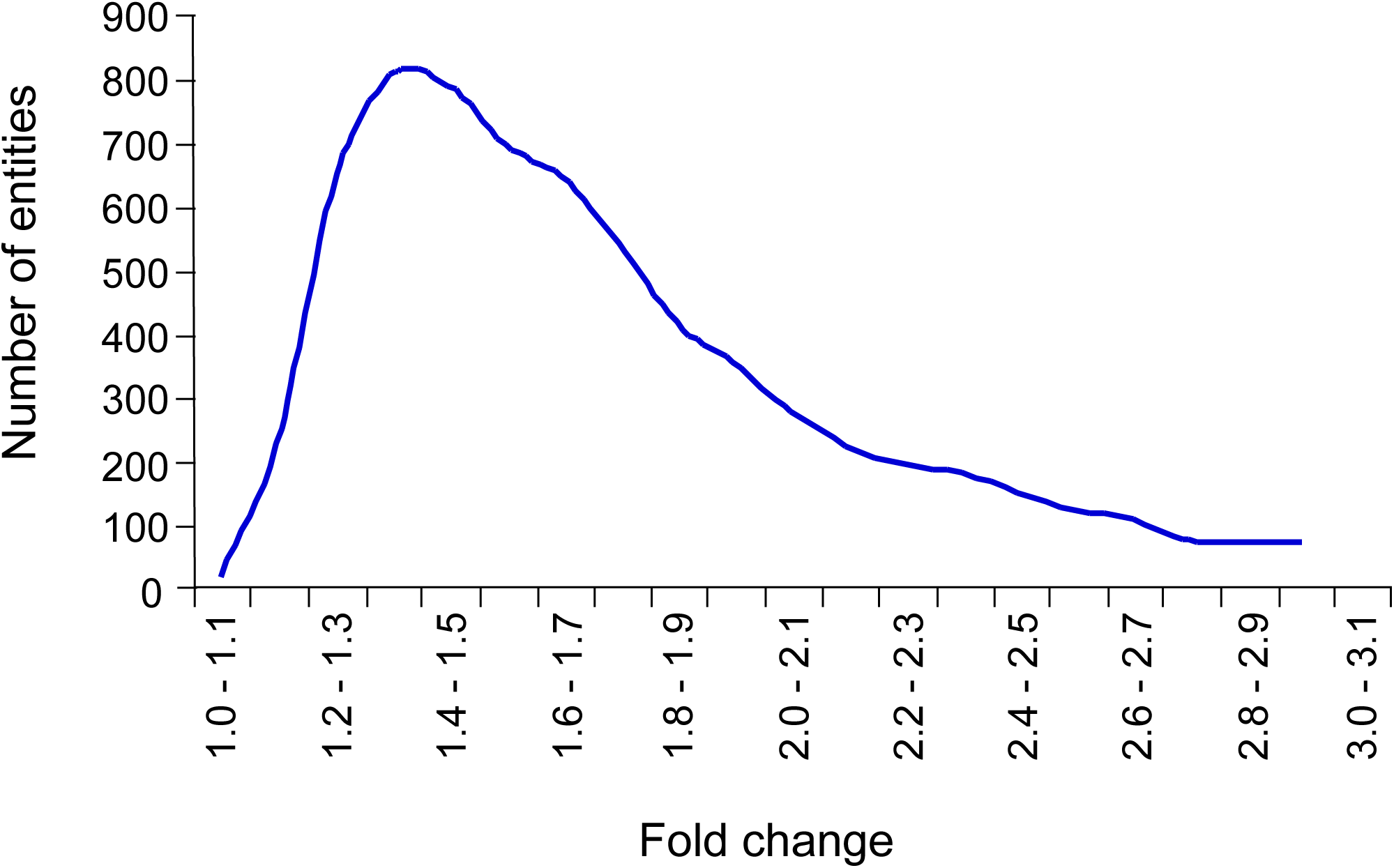
Allocation of differentially expressed genes in cultured cells relative to primary chondrocytes. A total of 14122 entities passed the criterion “present flag” in 13 out of 13 samples. In a second step 7256 entities were identified with a satisfying P-value cutoff 0.05. The highest number of genes with expression levels that were changed in chondrocytes under cultured conditions relative to primary chondrocytes were identified between 1.3- and 1.4-fold.

We then sought to identify major biological pathways in the temporal trajectories of expression in cultured cells. Because pathway analysis is more robust with larger gene lists, we ranked the 12078 entities detected at any stage in culture by initial expression level (day 2), divided them into quartiles, and built just three clusters for the predominant temporal trajectories in each quartile (Figure 7). Somewhat expected for genes with lowest initial expression (quartile 1), only increases in expression over time were detected, while in quartile 4 with the highest expression levels, most changes were reductions of expression over the course of culture. Furthermore, color coding in Figure 6 makes apparent that the vast majority of transcripts (7540, i.e. 62.4% of 12078) remain unchanged over the course of culture, regardless of expression level.

**Legend to Figure 7.**
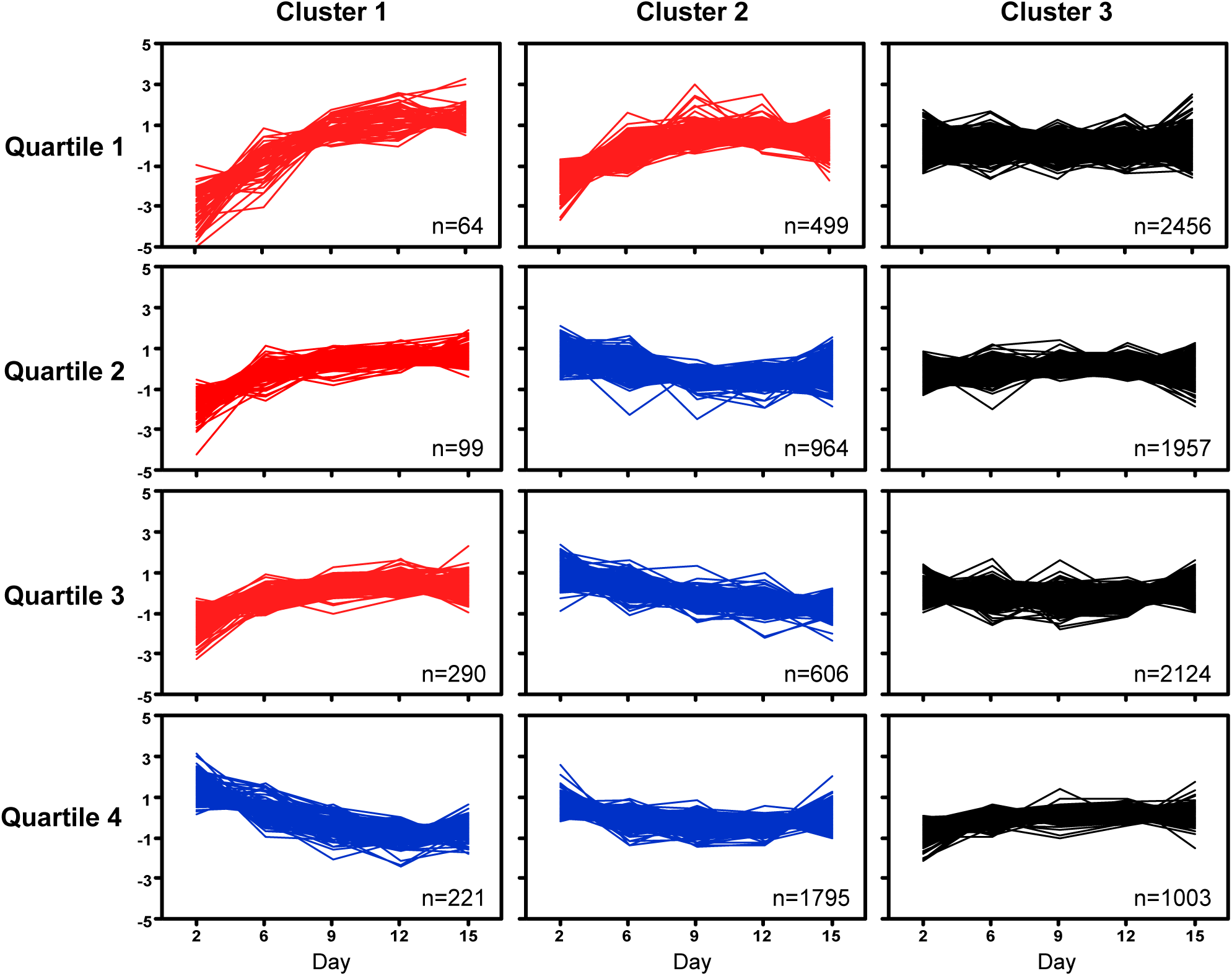
Differential gene expression in chondrocytes over time. K-Means clustering algorithm was used to visualize the different expression profiles. Intensity signals of 13078 Affymetrix probe IDs (intensity signal ≥ 500 in at least 1 out of 5 time points) were sorted ascending based on day 2 and separated into quartiles, then median normalized and log2 transformed. K-Means clustering was performed using JMP Genomics, where 3 clusters per quartile were built (n=number of probe IDs per cluster). The Figure depicts the median-normalized trajectory for each gene, with clusters ordered by trajectory pattern over time (red: increasing from initial expression level; blue: decreasing from initial expression level; black: no major change in expression level).

Pathway analyses (DAVID) revealed that clusters with constant expression (all black panels in Figure 7) were enriched for transcripts encoding proteins with roles in RNA transcription, protein synthesis and protein transport, cell cycle and mitosis, and mitochondrial function (enrichments factors ranging between 8.53 and 25.83). Thus, transcripts encoding basic cellular processes remain unchanged over the course of culture, although particular functions appear linked to expression level: overall, genes encoding transcription factors showed low expression (Q1), while genes with mitochondrial function exhibited higher expression (Q4). Additional functional categories with constant expression included ATP-synthesis, TCA-cycle, chaperones and unfolded protein response, ribosome and endoplasmatic reticulum, RNA processing/splicing, and DNA repair (in order of descending expression levels).

Among the genes with changing expression (4538, i.e. 37.6% of 12078), the largest group is comprised of transcripts that decrease over time (3586, i.e. 79% of 4538; blue in Figure 7). Since decreases from initially low expression levels would be technically difficult to detect, declining trajectories appear only in quartiles 2 through 4. These transcripts were enriched for genes with roles in mitosis and DNA replication, RNA transcription and processing, ribosomal translation and ribosome biogenesis, and protein synthesis, consistent with declining chondrocyte proliferation as the cells mature in culture. Of transcription factor genes known to be expressed in cartilage, decreases were found for Cebp*β*, Cebp*ζ*, Ets1, Klf5, Klf10, Max, Myc, Sox6, Sox9, and Twist2. The phenotypes of knockout mouse mutants (as annotated in http://www.informatics.jax.org) for these genes have in common reduced cell proliferation, consistent with lower expression levels during the later, less proliferative^10^, stages of the chondrocyte cultures.

Genes with increasing expression levels comprised the smallest group, with 952 genes (i.e. 7.88% of 12080, or 21% of 4538 entities with expression changes; red in Figure 7), suggesting that activation of transcription is possibly harder to achieve in culture than reduced expression. Transcripts with greatest increases (in Q1 and Q2) were enriched for components of cartilage and extracellular matrix, such as Collagens (1a2, 3a1, 5a1, 8a1, and 12a1), Fibromodulin, Matrilin, Comp, Decorin, Integrin-binding-sialoprotein, Thrombospondin2, Tenascin, and Versican, for examples. Also, secreted molecules with roles in cell adhesion and cartilage and bone development were prominent, such as Igf1 and Igf-binding-protein4, Cadherin11, Galectin3, Fibulin, Fibroblast-activation-protein, Gdf10, Gremlin, Mmmp13, Mmp14, Midkine, PdgfC.

Genes with increasing expression levels also included many transcription factors known to play a role in chondrocyte differentiation, such as Ahr, Atrx, Cepb*α*, Creb3-like1, Fos, Foxn3, Hoxa9, Hoxb3, Hoxd8, Maf, Mxd4, Medf2c, Nr2f2, Nr4a1, Pbx1, Pitx2, Prrx2, Runx1, Sox4, Sp7, Stat1, Stat3, and Twist1 (For a complete list of transcription factors see Supplemental Table 3). The parallel trajectories of increasing expression over time in culture suggest that these regulators could play direct roles in the activation of transcription of genes encoding the secreted and extracellular matrix components. Furthermore, the limited number of regulators and potential targets provides evidence that the chondrocyte maturation program is orchestrated by highly specific molecular mechanisms.

## DISCUSSION

Here, we sought to investigate the molecular mechanisms underlying chondrocyte differentiation and maturation in a defined culture system that employs primary cells from ribs of normal fetal mice. Our high-density culture system recapitulates the *in vivo* sequence of chondrocyte maturation, by virtue of extracellular matrix production, cellular hypertrophy and, at late stages of culture, mineralization. The results from our expression profiling confirm the involvement of molecules that are well-known from prior studies as regulators and components of cartilage maturation, including transcription factors (Sox and Hox, for examples) and extracellular matrix molecules (Collagens, Matrilin, Comp, Tenascin and others). Furthermore, our unbiased genome-wide survey revealed additional pathways for which extensive literature is currently not available. For example, the parallel temporal trajectories of transcription factor and extracellular matrix genes provide strong plausibility for the notion that these transcription factors are direct regulators of extracellular matrix production. Conversely, inverse temporal profiles for transcription factors and potential targets would be consistent with negative regulation, i.e. silencing or repression, of target transcription. Thus, our results raise new testable hypotheses about molecular mechanism in cartilage production and maturation.

The second major outcome is the finding that, even within two days, cells in culture had noticeably altered gene expression repertoires compared to freshly isolated chondrocytes. This partly reflects the disappearance (probably through cell death) of some initially contaminating non-cartilage cell types, such as blood and muscle cells. But we also found that expression levels of some known chondrocyte-specific genes were higher in fresh cells compared to day 2 in culture, such as Chondroadherin (Chad), Chondrolectin (Chodl), Glycosyltransferase25-domain-containing-2 (Glt25d2), Cartilage-intermediate-layer-protein-2 (Clip2), TenascinXB, Fos, and Collagen9a2. Thus, transcript levels re-calibrated as cells accommodated to the altered environment and significantly increased expression of genes with roles in cell differentiation, adhesion and extracellular matrix production, especially of well-known cartilage components Matrilin, Comp, and Proteoglycans. Thus, despite the loss of expression for many genes from the time of isolation from the animal, high-density culture of rib-derived chondrocytes supports their maturation, and production of copious cartilage material. Our profiling studies also revealed expression of genes whose function in chondrocytes was previously unknown. Based on their temporal profile of elevated transcription at later stages of chondrocyte differentiation, it is conceivable that these genes are involved in maintenance of the differentiated chondrocyte cell fate.

Overall, rib-derived cells reproduce the steps in cartilage maturation that were also observed in micromass cultures when embryonic limb mesenchyme^9, 19^ was used as a source. Yet, in comparison of our results to microarray studies from micromass cultures^19^, some differences are notable: At day 3 of culture, James et al., 2005^19^ detected strong expression of myocyte genes that decreased only at later stages of culture. In contrast, such transcripts are already reduced by day 2 of our rib chondrocyte cultures. This indicates that either some limb-derived cells differentiate into muscle in culture, or if already present in the embryonic limbs at the time of isolation, they are slower to die off. The results also could reflect a more advanced stage of chondrocyte differentiation in primary rib cartilage cells from compared to limb mesenchyme. Consistent with this interpretation, we observe increases of several markers of chondrocyte maturity towards very high expression levels at later stages of rib-derived cultures that remain moderate, or even decrease, in the limb micromass cultures, such as for example for Adamts2 and Adam23, and Otor/CD-RAP and Comp, respectively.

In a later report, James et al., 2010 compared their expression profiles from micromass cultures to fresh tissue from different tibial regions of gestational day 15.5 mouse embryos^20^. In analogy to our results, they find considerable differences between cultured cells and fresh samples, with enrichment in the cultures of genes with roles in cellular physiology, mitochondrial activity, protein transport and extracellular matrix remodeling. A caveat here is that these published reports differ from our parameters for microarray data analysis, clustering and gene set enrichment methods, as well as in annotation approaches and the underlying database versions.

We therefore performed a direct comparison, by re-processing (based upon Affymetrix probe identifiers) and re-analyzing all datasets with uniform methodology. To visualize relationships and identify factors accounting for possible differences, we conducted Principal Component Analysis (Figure 8; Beier laboratory data in red, our data in blue). Panel A highlights that Principal Component 1 (PC1) accounts for almost 40% of the overall variation between samples. Visual inspection in three-dimensional space indicated that PC1 was driven by the spread of the time course series from both groups, implicating time as the major factor. Interestingly, the analysis separated the results from both laboratories along PC2: all red symbols, regardless of shape, fell between -0.2 and -0.1 on the PC2 axis, while all blue symbols located between 0.1 and 0.2. Thus, PC2 –the difference between laboratories– accounts for 23.5% of the variation, highlighting approach-specific, experimenter-specific or other laboratory-specific factors. Any remaining components accounted for less than 10% of the variation, as indicated by 9.3% for PC3. Removing PC2 from the plot (Panel B) did not change the spread of samples along PC1 –as expected–, and all samples also occupied the same relative position in the PC3 dimension. PC4 provides 6.5% of the variation, but the major distinctions between the timecourse data, and the primary tissues, for either laboratory do not change substantially. Notably, the biological replicates for primary tissue cluster very closely together (Panels A and B), highlighting consistency of dissection and sample preparation in each laboratory. Not unexpectedly, once the laboratory-specific factors are removed, the location of rib chondrocytes (labeled RC in Figure 8) is close to growth plate Zone I samples dissected from embryonic tibiae, consistent with the presence of proliferating chondrocyte precursors in primary cartilage tissues.

**Legend to Figure 8.**
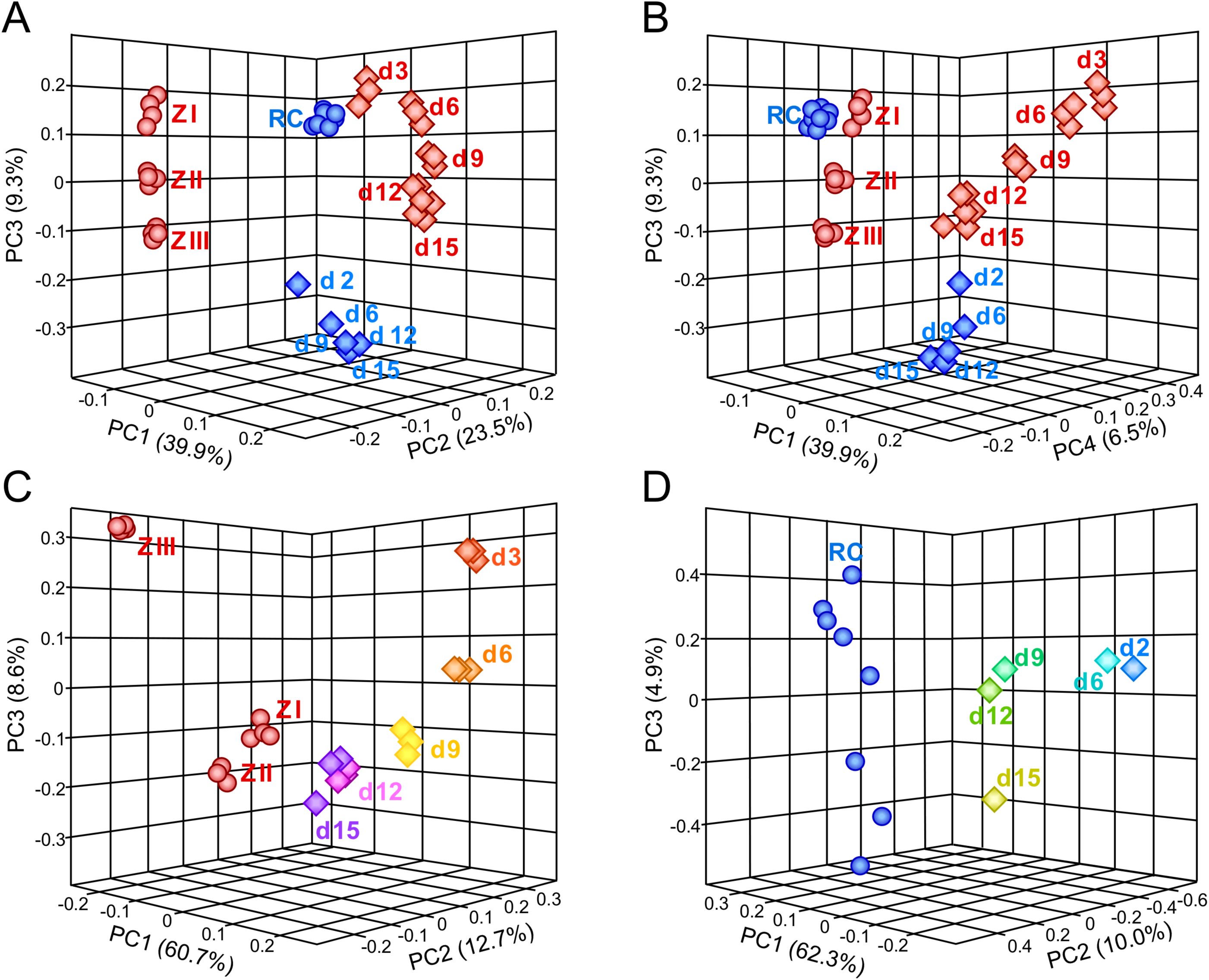
Comparison of independent timecourse experiments for chondrocyte differentiation. Principal Component Analyses were performed in JMP Genomics. Panels A and B depict the results for the combined dataset, Panels C and D depict the analysis results for the data from the Beier group (C) and the Kappen laboratory (D) separately. Note that scales are different in each graph. Microarray samples from the Beier lab are represented by red symbols, samples from the Kappen lab are in blue. Primary tissue is represented by spheres, for dissected E15.5 embryonic tibiae Zones I through III (in red, ZI - ZIII), and primary rib chondrocytes from dissected newborn ribs (in blue, RC). In Panels A and B, rectangles represent the samples from micromass cultures of embryonic limb mesenchyme (in red), and samples from high-density cultures of newborn rib chondrocytes (in blue). In Panels C and D the rectangles are colored by day of culture as labeled. PC = Principal Component.

Separate analyses confirmed time (PC1) as the major factor driving variability, with 60.7% and 62.3% weight (in Panels C and D, respectively). Such wide spread is not unexpected, as initially undifferentiated limb mesenchymal cells commit to the chondrocyte lineage only after several days in the micromass culture^19^. The data from our laboratory also spread along PC1, albeit not to the same extent, indicating that starting and differentiated cells in the high-density cultures are more similar to each other than in the micromass assay.

Further inspection revealed convergence of results from both laboratories for cultured cells over time, with d15 and d12 samples closer to each other than earlier time points. Multivariate correlation analyses for the 1000 highest expressed genes defined 105 probes as different by more than 2-fold between micromass and high-density cultures in the day 2-to-3 comparison, then decreasing to 46 probes at day 12, and 32 probes at day 15. Fold-changes of expression differences were also greater at early time points. The small number of genes accounting for final differences highlights the overall strong concordance of temporal expression profiles from both culture models.

In conclusion, extensive gene expression profiling of cartilage differentiation, as reported here and by others^19, 20^, confirmed -notwithstanding considerable cellular adaptation to the *in vitro* conditions- the validity of culture models for cartilage formation. We also showcase opportunities for higher-level systems analyses that can further elucidate temporal mechanisms and identify potentially novel regulators and pathways in cartilage differentiation and maintenance of the mature chondrocyte phenotype.

## ACKNOWLEDGEMENTS

We wish to thank Ms. Cathy Huey for administrative assistance and Susan Newman (PBRC Genomics Core) for expert support of the bioinformatics analyses. Funding was provided in parts from a grant by the Aircast Foundation, National Institutes of Health grants R21-AR052731 and R21-DE014523, and the Peggy M. Pennington Cole Endowed Chair at Pennington Biomedical Research Center. The Genomics Core Facility at PBRC is supported in parts by Center for Biomedical Research Excellence P30GM118430 and Nutrition and Obesity Research Center P30DK072476 grants from the National Institutes of Health.

## DECLARATIONS

### Author Contributions

Aimee Limpach performed cell culture experiments, RNA isolation and RT-PCR assays, and microarray analyses; Claudia Kruger performed RT-PCR assays, microarray data analysis, and bioinformatics analyses, the Microarray Facility at University of Nebraska Medical Center conducted the microarray assays; Claudia Kappen performed pathway analyses and wrote the manuscript. All authors read and approved the manuscript before submission.

### Data availability statement

The datasets supporting the conclusions of this article are included within the article and the Supplemental Files. The primary microarray data have been deposited to GEO (GSE102292) and will be available upon publication of this manuscript.

### Ethical approval and informed consent

All animal experiments were conducted in accordance with applicable state and federal regulations (https://grants.nih.gov/grants/olaw/guide-for-the-care-and-use-of-laboratory-animals.pdf). The most recent Institutional Animal Care and Use Committee approval relevant to this report was issued in December 2006, PBRC protocol #442.

**Legend to Supplemental Table 1.**

**Investigated genes by qRT-PCR: primer sequences and amplification efficiency.**

Sequences were taken from the ENSEMBL Genome Browser. Where possible, the amplicon span an exon-exon junction. The amplification rate was calculated with the formula: *Δ*Rn_cycle (n) / *Δ*Rn_cycle (n-1).

**Legend to Supplemental Table 2.**

**Functional classification of genes altered in response to culture time.**

Enrichment analysis was performed on 12 clusters graphed in Figure 7 using DAVID functional annotation clustering based on SP_PIR_Keywords and GOTERM_FAT categories.

**Legend to Supplemental Table 3.**

**Changes in transcription factor expression over the time course in culture.**

Gene symbols (1675) of mouse transcription factors were downloaded from the Riken Transcription Factor Database (http://genome.gsc.riken.jp/TFdb) and compared to gene lists obtained after K-Means clustering. Total number of probe IDs and signal intensity range per quartile are noted. Ingenuity Pathway Analysis results for potential targets of transcription factors are given in separate worksheets for each comparison.

**Legend to Supplemental Table 4.**

**Expression of transcription factor genes in cultured chondrocytes by expression level**

Transcription factor genes within each cluster were annotated according to the Riken Transcription Factor Database (http://genome.gsc.riken.jp/TFdb).

